# Separating anorexia -dependent and -independent effects in cancer cachexia

**DOI:** 10.1101/2025.04.24.650538

**Authors:** Yanshan Liang, Young-Yon Kwon, Sheng Hui

## Abstract

Cancer cachexia is characterized by unintentional weight loss and wasting away of fat and muscle tissues. Anorexia, or reduced food intake, is often implicated as a contributor to the negative energy balance in this condition. However, to what extent anorexia alone accounts for body weight loss and wasting of different tissues, and whether anorexia is responsible for other cachectic phenotypes such as physical performance impairment remains insufficiently characterized in preclinical models and patients. In this study, we demonstrate the critical need to address these questions in cancer cachexia research. Using the colon carcinoma 26 (C26) model of cancer cachexia as an example, we systematically determined how much each of the key phenotypes of cancer cachexia is driven by anorexia. Anorexia was the predominant driver for body weight loss, adipose tissue wasting, and muscle wasting, strikingly suggesting the lack of other mechanisms for causing these phenotypes in this model. In contrast, anorexia had no impact on physical performance, pointing to the existence of anorexia-independent mechanisms in causing fatigue. Thus, for a given preclinical model or patient group, anorexia can be the main cause for certain cachectic phenotypes and play no role in causing other cachectic phenotypes. Discriminating between anorexia-mediated and independent effects is essential for guiding research focus and ultimately unraveling the causal pathways of cancer cachexia.

## Introduction

Cancer cachexia is an involuntary wasting condition prevalent in cancer patients, contributing to poor quality of life and reduced survival [1,2]. However, the etiology of the disease remains to be fully understood, and no effective treatment is currently available [3]. One challenge in establishing the causal pathways of cancer cachexia is to distinguish causality from mere association, as evidenced in our previous work showing that the widely assumed cachectic factor IL-6 likely only associates with cancer cachexia [4]. Another often-cited factor contributing to cancer cachexia is anorexia. However, to what extent anorexia contributes to the body weight loss and also to other phenotypes of cancer cachexia, such as loss of fat mass and muscle mass, impairment of physical performance, and alteration of systemic metabolism, has been rarely rigorously assessed in preclinical models or human patients. This is a critical question because the answer dictates the focus of downstream investigations. For example, food intake reduction is known to cause reduced muscle mass even in healthy animals [5,6]. If anorexia alone accounts for most muscle mass loss in cancer cachexia, research should be focused on unraveling mechanisms causing anorexia rather than some active mechanisms of muscle wasting. The same question can be posed for other phenotypes of cancer cachexia, such as adipose tissue wasting and metabolic alterations, both of which are heavily influenced by food intake [7,8]. Thus, in this study, we focus on the causality of food intake reduction in different cachectic phenotypes and demonstrate the importance of separating anorexia-dependent and -independent effects in cancer cachexia research.

Among previous studies of cancer cachexia, a marginal fraction of them controlled for food intake. While the lack of food intake control in patient research is likely due to practical infeasibility, we were only able to find on the order of 10 preclinical studies that used a pair-feeding strategy (matching daily food intake between cachectic and non-cachectic groups) [9–14] and a couple that employed a caloric restriction strategy (matching total food intake between cachectic and non-cachectic groups) [15,16]. These studies focused on one or two specific phenotypes such as body weight or adipose tissue wasting, and missed key phenotypes such as physical performance. Thus, a study is needed to systematically delineate the causal roles of anorexia in all key phenotypes of cancer cachexia.

Here, we systematically measured how reduced food intake contributes to cachexia phenotypes and metabolic changes in the syngeneic murine colon 26 carcinoma (C26) cancer cachexia model [17] using an isocaloric feeding strategy. Our findings revealed that anorexia is a primary driver for body weight loss, fat mass loss, and muscle protein loss in this model, suggesting the lack of additional mechanisms for these cachectic phenotypes. In contrast, anorexia does not affect physical performance, highlighting anorexia-independent cachectic pathways. Additionally, anorexia is responsible for key alterations of systemic metabolism, including reduced circulating glucose and increased ketone body levels, suggesting no need to invoke additional mechanisms to explain these metabolic changes. Therefore, distinguishing anorexia-induced effects from other cachectic symptoms is crucial for guiding research efforts to identify the causal pathways of cancer cachexia.

## Results

### Anorexia accounts for almost all body weight loss and body composition changes in the C26 model

We first aimed to investigate the contribution of anorexia to body weight loss, a primary phenotype of cancer cachexia, in the C26 model. The C26 model was set up by subcutaneously injecting the C26 cells into the syngeneic CD2F1 mice[4,17]. Two weeks post-tumor inoculation, these mice exhibited over 15% body weight loss manifested by fat and lean mass reductions. We refer to this group as the cachectic C26 (cxC26) group. To control for effects due to tumor growth, we included a mouse group using a non-cachectic C26 cell line (ncxC26), which maintained body weight and composition despite similar or bigger tumor size (Supplementary Fig. S1).

The cxC26 mice showed clear anorexia (Fig. 1a). Consistent with previous observations [18–20], these mice produced a large amount of masticated food debris (Supplementary Data Fig. S2). For accurate food intake measurement, mice were single-housed in cages with paper bedding, which allows reliable separation of food debris and bedding materials. To determine the extent to which anorexia contributes to body weight loss, we initially employed a pair-feeding approach by providing a group of ncxC26 mice with a single portion of food equal to the amount consumed by cxC26 cachectic mice ad lib. The pair-fed ncxC26 mice showed similar body weight loss as the cxC26 mice under comparable tumor burden (Supplementary Fig. S1a, c). Fat and lean mass reductions were consistently comparable between the pair-fed ncxC26 and cxC26 mice, suggesting a predominant role for anorexia in body weight loss and composition (Supplementary Fig. S1b). A potential caveat in the pair-feeding experiment is that pair feeding alters the diurnal food intake patterns and subjects the animals to a fixed feeding schedule, which can induce caloric deprivation and lead to different metabolic and physiological responses in freely-fed animals [21,22]. Thus, to minimize potential differences in feeding behavior between the two groups, we adopted an isocaloric feeding approach in which both ncxC26 and cxC26 mice were provided daily the same amount of food matching the food intake of a group of cxC26 mice ad lib as shown in Fig. 1a. Similar to the pair-feeding experiment, the body weight curves were in close match between the isocaloric-fed ncxC26 and cxC26 groups (Fig. 1b-c). Furthermore, the extent of both lean mass loss and fat mass loss was also the same between the two isocaloric-fed groups (Fig. 1d). Importantly, there was no difference in tumor size between the two groups (Fig. 1e). These results showed that anorexia accounts for almost all of the three key cachectic phenotypes: body weight loss, fat mass loss, and lean mass loss.

**Figure 1.**
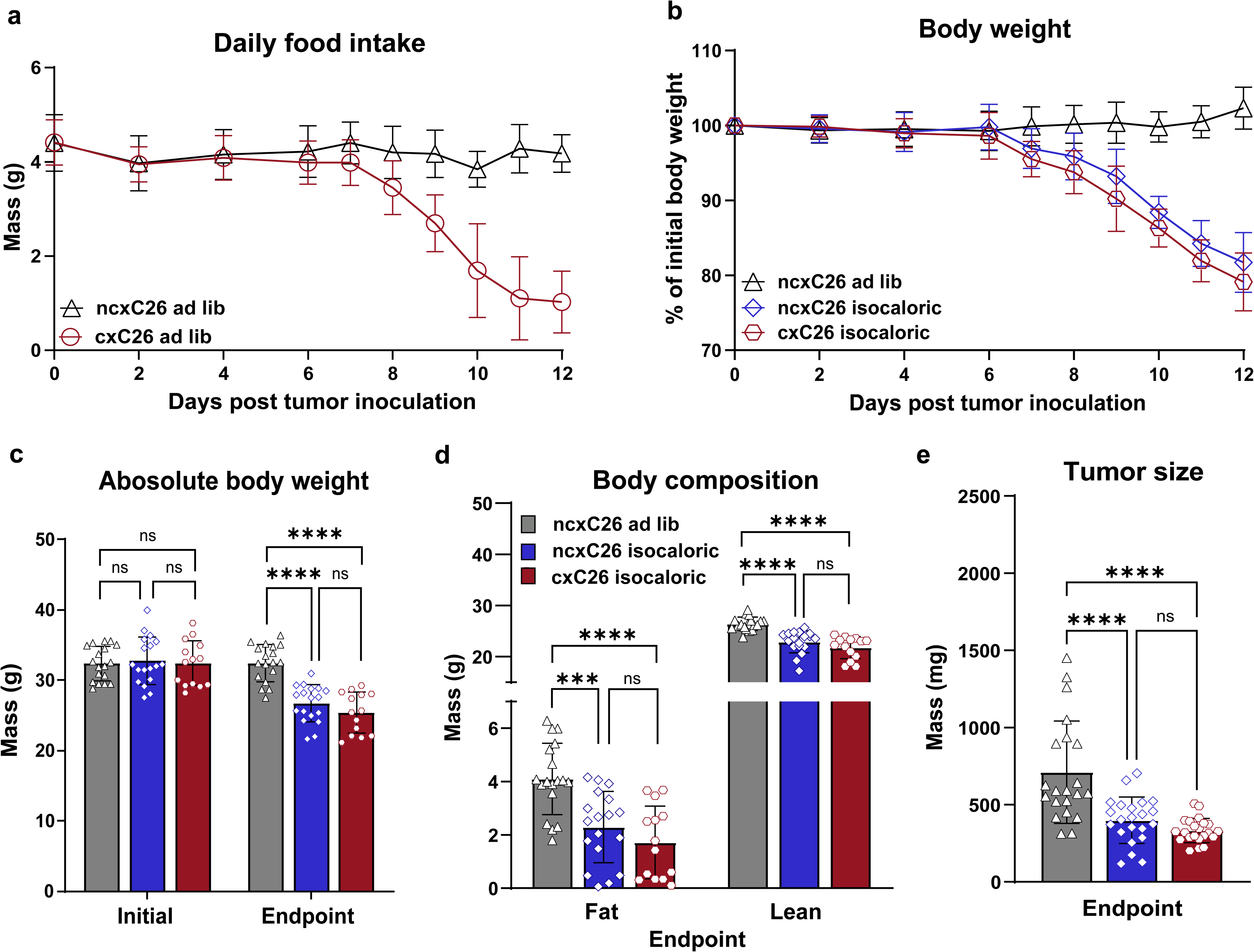
Anorexia is the main driver of body weight loss and body composition changes. CD2F1 male mice were inoculated with 1×10^6^ ncxC26 or cxC26 cells subcutaneously. **(a)** Food intake of ncxC26 and cxC26 mice ad lib. **(b)** Percentage change in body weight, **(c)** absolute initial and endpoint body weight, (**d**) body composition measured using EchoMRI at the endpoint, and **(e)** tumor size at the endpoint, for ncxC26 ad lib, isocaloric-fed ncxC26, and isocaloric-fed cxC26 groups. **(a)** n=16 for ncxC26 ad lib and n=16 for cxC26 ad lib. **(b)** n=20 for ncxC26 ad lib, n=20 for isocaloric-fed ncxC26, n=15 for isocaloric-fed cxC26. **(c-d)** n=18 for ncxC26 ad lib, n=18 for isocaloric-fed ncxC26, n=14 for isocaloric-fed cxC26. **(e)** n=21 for ncxC26 ad lib, n=22 for isocaloric-fed ncxC26, n=22 for isocaloric-fed cxC26. All data are shown as the mean ± s.d. Significance of the differences: ns not significant, *P < 0.05, **P < 0.01, ***P < 0.001, **** P<0.0001 between groups by one-way ANOVA followed by Tukey post hoc comparison.

### Neither energy expenditure nor fecal energy excretion contributes to body weight loss

Our observation that anorexia is the sole driver for body weight loss implies a limited role for energy expenditure (EE) in body weight loss, which is often cited to be concomitant with cancer cachexia [23,24]. To test this, we measured the EE using a metabolic cage system for the isocaloric-fed ncxC26 and cxC26 mice. With identical caloric intake, the time-course EE data in cxC26 mice closely followed that of ncxC26 mice, which was lower than the ncxC26 mice ad lib (Fig. 2a). The lack of difference in EE between the two isocaloric-fed groups was also evident in the cumulative EE data (Fig. 2b). The metabolic cage data further showed no difference between the two groups in the respiratory exchange ratio (RER), indicating their similar substrate utilization at the whole-body level (Fig. 2c-d).

**Figure 2.**
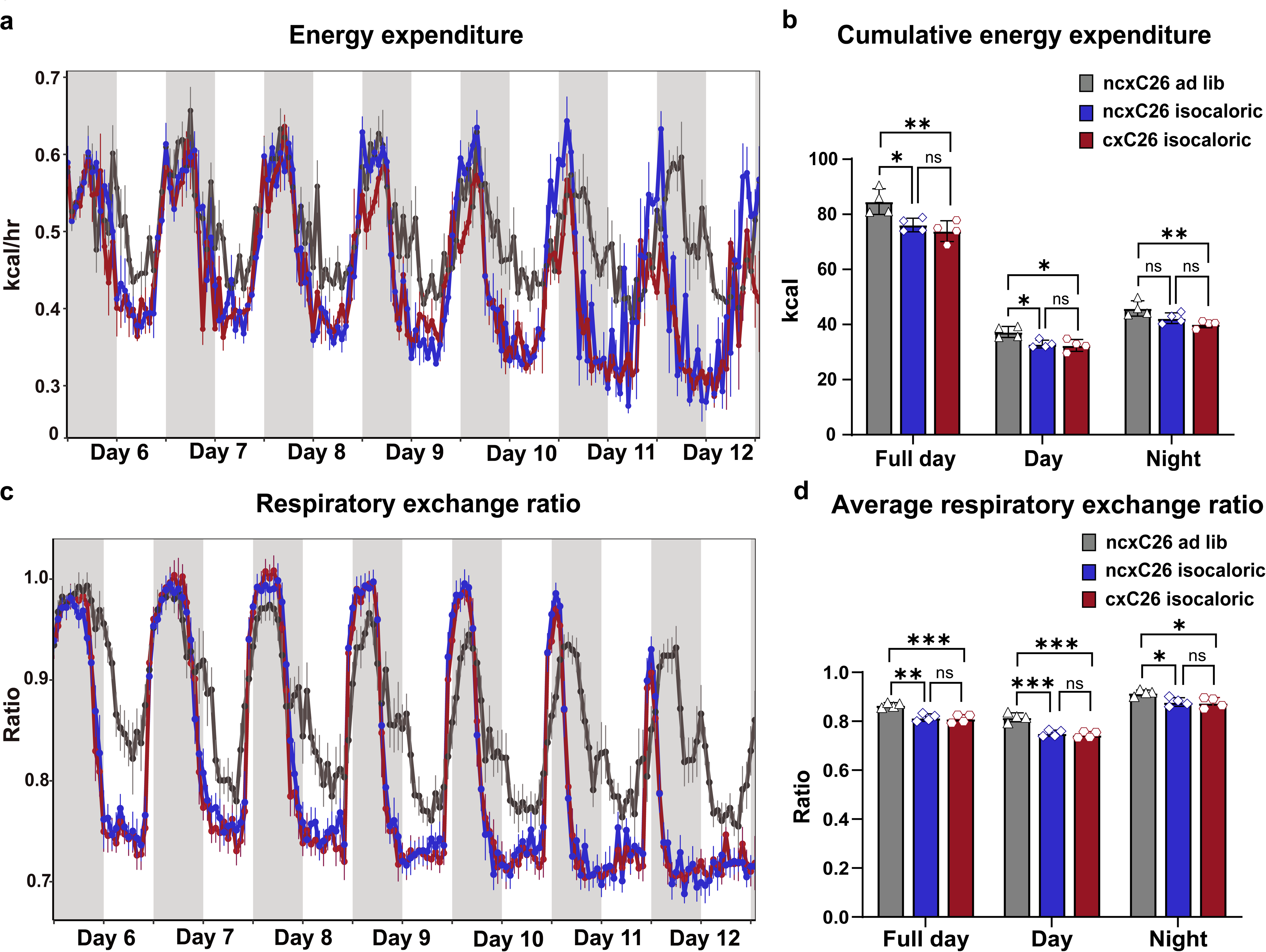
Energy expenditure does not contribute to body weight loss. **(a)** Hourly energy expenditure from day 6 to day 12 post-inoculation. **(b)** Cumulative energy expenditure during full day, daytime, and nighttime. **(c)** Hourly respiratory exchange ratio from day 6 to day 12 post-inoculation. **(d)** Average respiratory exchange ratio. **(a-d)** n=4 per group. **(b and d)** are shown as the mean ± s.d. Significance of the differences: ns not significant, *P < 0.05, **P < 0.01, ***P < 0.001, **** P < 0.0001 between groups by one-way ANOVA followed by Tukey post hoc comparison.

Another important variable in the energy balance equation is energy excretion, which can be affected by possible malabsorption complications [25,26] in cancer cachexia. To determine its role in energy balance and body weight loss, we collected all feces produced within the 24 hours before the endpoint from the three groups. We then measured fecal calories using bomb calorimetry. Our results indicated that the two isocaloric-fed groups had similar fecal production though significantly less than the ncxC26 mice ad lib (Fig. 3a). As the fecal energy density was not changed across the groups (Fig. 3b), there was reduced but identical total fecal energy excretion in the two isocaloric-fed groups (Fig. 3c). Together, these results demonstrated that neither energy expenditure nor fecal energy excretion contributes to body weight loss; instead, anorexia accounts for almost all body weight loss as well as body composition in the C26 model.

**Figure 3.**
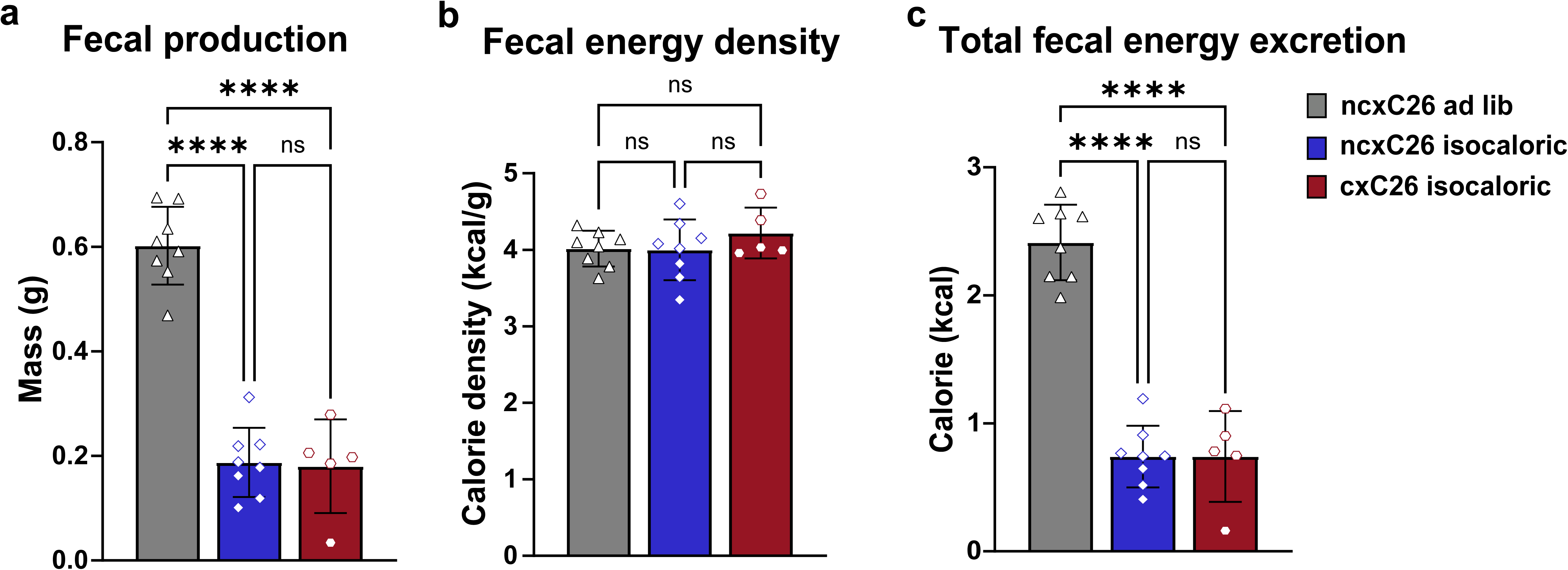
Energy excretion does not contribute to body weight loss. Feces produced by mice within 24 hours before euthanasia were collected. **(a)** Fecal production, **(b)** Fecal energy density **(c)** Total fecal energy excretion. **(a-c)** n=8 for ncxC26 ad lib, n=8 for isocaloric-fed ncxC26, and n=5 for isocaloric-fed cxC26. All data are shown as the mean ± s.d. Significance of the differences: ns not significant, *P < 0.05, **P < 0.01, ***P < 0.001, **** P < 0.0001 between groups by one-way ANOVA followed by Tukey post hoc comparison.

### Anorexia is the main driver for muscle protein mass loss in cancer cachexia

Skeletal muscle wasting is a key hallmark of cancer cachexia, and the tumor-secreted factors have been reported to have direct effects on muscle [27,28]. Our body composition data, however, suggested that the muscle wasting could be simply due to anorexia. To investigate further the effects of anorexia on skeletal muscle mass, at the endpoint, we dissected and weighed the lower limb muscles, including gastrocnemius, soleus, tibialis anterior (TA), and extensor digitorum longus (EDL). Consistent with the lean mass result, reduced food intake for the ncxC26 group resulted in generally reduced muscle mass (comparing the isocaloric-fed ncxC26 group to the ncxC26 ad lib group), albeit to slightly less degree than the cxC26 group in gastrocnemius and soleus (Fig. 4a). The muscle mass change was not due to glycogen mass change because of low muscle glycogen content (<1%) (Fig. 4b). As proteins are the main component of muscle tissue, we next assessed the protein content of the gastrocnemius muscle. We observed no significant difference in the protein content of gastrocnemius muscle between isocaloric-fed cxC26 and ncxC26 mice (Fig. 4c), indicating similar magnitude of muscle protein mass loss in the two groups (Fig. 4d). This finding was corroborated by the gene expression levels of muscle atrophy biomarkers *Murf1 and Atrogin1* (Fig. 4e), which were elevated in the ncxC26 group to the same level as in the cxC26 group. Altogether, these results showed that it is reduced food intake, not some specific cachectic factor, that is the main driver for muscle protein wasting in the C26 model.

**Figure 4.**
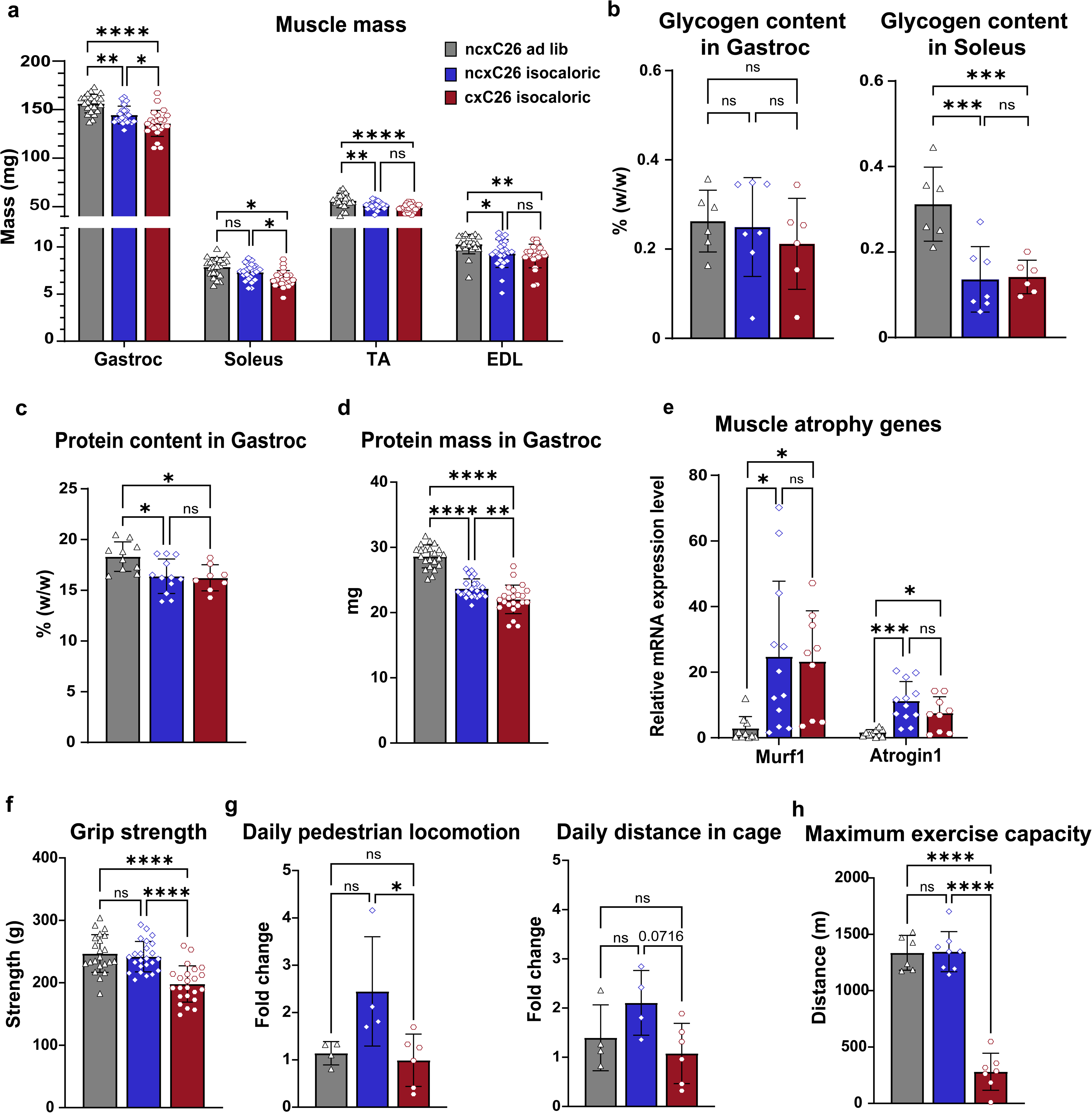
Food intake reduction induces muscle protein wasting but not fatigue in cancer cachexia. **(a)** Muscle mass. **(b)** Muscle glycogen content. **(c)** Muscle protein content. **(d)** Total protein mass in gastrocnemius calculated by multiplying averaged protein content with muscle mass. (**e)** mRNA expression levels of atrophy markers in gastrocnemius. **(f)** Grip strength. **(g)** Changes in daily pedestrian locomotion and distance in cages comparing D12 to D0. **(h)** Maximum exercise capacity (running on a treadmill until exhaustion). **(a, d)** n=23 for ncxC26 ad lib, n=24 for isocaloric-fed ncxC26, n=28 for isocaloric-fed cxC26. **(b)** n=6 for ncxC26 ad lib, n=7 for isocaloric-fed ncxC26, n=6 for isocaloric-fed cxC26. **(c)** n=10 for ncxC26 ad lib, n=12 for isocaloric-fed ncxC26, n=8 for isocaloric-fed cxC26. **(e)** n=10 for ncxC26 ad lib, n=12 for isocaloric-fed ncxC26, n=9 for isocaloric-fed cxC26. **(f)** n=22 for ncxC26 ad lib, n=23 for isocaloric-fed ncxC26, n=23 for isocaloric-fed cxC26. **(g)** n=4 for ncxC26 ad lib, n=4 for isocaloric-fed ncxC26, n=6 for isocaloric-fed cxC26. **(h)** n=6 for ncxC26 ad lib, n=8 for isocaloric-fed ncxC26, n=7 for isocaloric-fed cxC26. All data are shown as the mean ± s.d. Significance of the differences: ns not significant, *P < 0.05, **P < 0.01, ***P < 0.001, **** P < 0.0001 between groups by one-way ANOVA followed by Tukey post hoc comparison.

### Physical performance is independent of anorexia-induced muscle mass loss in cancer cachexia

To continue assessing the effects of anorexia on key phenotypes in cancer cachexia, we next determined the impact of anorexia on physical performance. For this, we first measured grip strength and found surprisingly that compared to the ncxC26 ad lib group, despite reduced muscle mass, the isocaloric-fed ncxC26 group did not show a decrease in grip strength (Fig. 4e). In contrast, the isocaloric-fed cxC26 group showed clear reduced grip strength (Fig. 4e). Grip strength impairment has been reported to be closely associated with fatigue [29–31]. To subject this finding to further testing, we next measured physical activity, which is well related to fatigue [32] and used for fatigue evaluation [33]. Both daily pedestrian locomotion and daily distance covered in the cage showed increased activity in the isocaloric-fed ncxC26 group, reflecting a foraging behavior under starvation, but not in the isocaloric-fed cxC26 group (Fig. 4f). This observation, in line with the grip strength results, suggests that anorexia has a limited role in contributing to fatigue. Finally, we performed an additional assay to evaluate fatigue-like behavior [34]. In this test, mice were forced to run on a treadmill until exhaustion to assess their maximal endurance performance. Again, consistent with the grip strength and the physical activity data, the isocaloric-fed cxC26 group showed a substantial reduction in performance while the isocaloric-fed ncxC26 group did not (Fig. 4g). Together, these results pointed to limited role of anorexia in contributing to fatigue.

The independence of fatigue on muscle mass is a striking result. A potential caveat for this conclusion is that cachectic animals exhibited slightly but significantly smaller muscle mass compared to their isocaloric-fed non-cachectic counterparts (Fig. 4a). Thus, to eliminate this potential caveat and more rigorously test our finding, we subjected a group of ncxC26 mice to a severe caloric restriction, reducing their food intake by 30% relative to that of the cxC26 mice ad lib starting from day 7 post-tumor injection (Fig. 5a). With this approach, the body weight, lean and fat mass, and muscle mass of the caloric-restricted ncxC26 mice were aligned with, and in some cases slightly lower than, those of the cxC26 mice ad lib (Fig. 5b-d). We employed the grip strength assay for fatigue evaluation and observed that the grip strength of ncxC26 caloric-restricted mice was comparable to that of the freely fed ncxC26 mice despite substantial food intake reduction (Fig. 5e). As the caloric-restricted ncxC26 group now unambiguously had matched muscle mass with the cxC26 group, their largely intact physical performance clearly demonstrated that fatigue is independent of anorexia-induced muscle mass loss.

**Figure 5.**
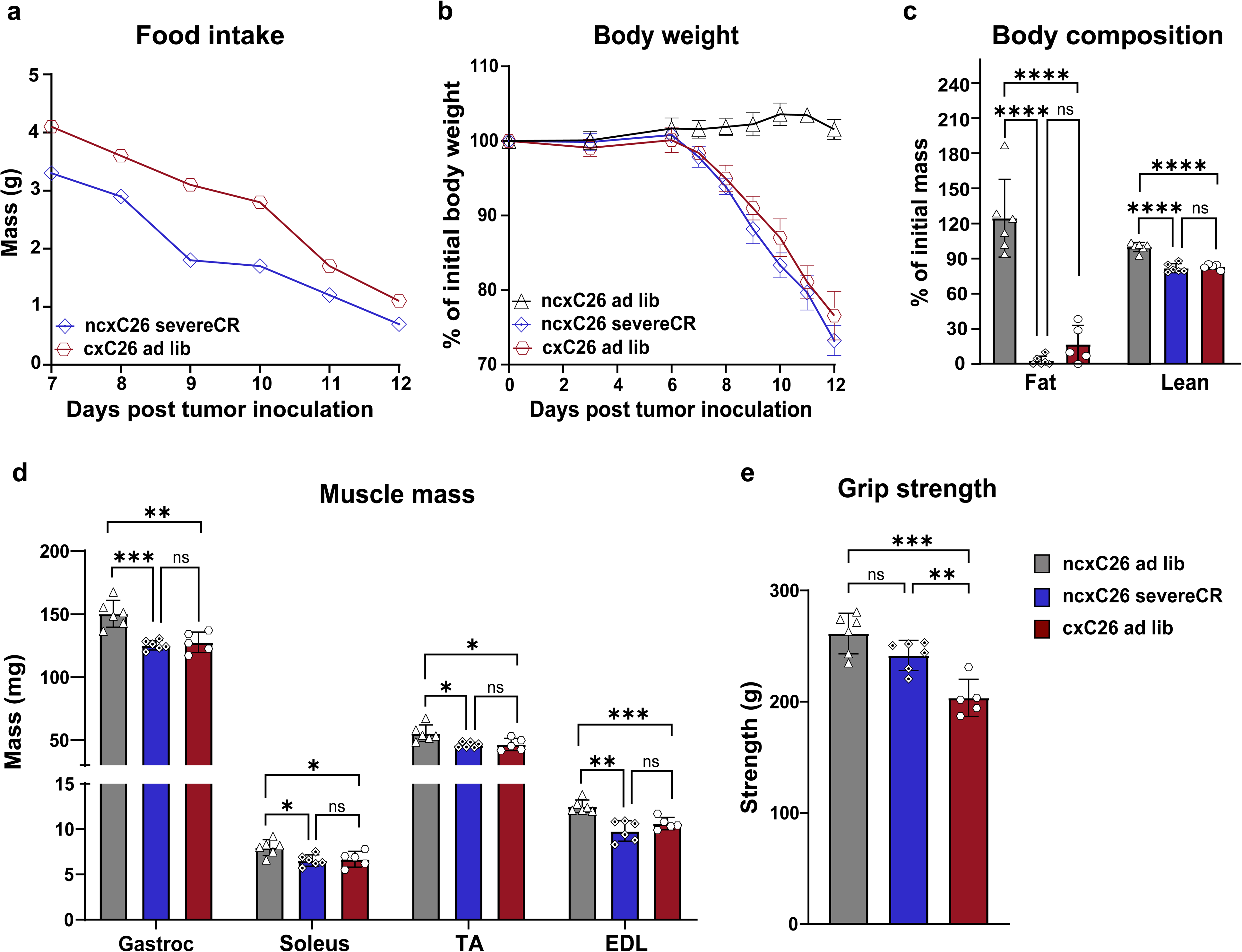
Fatigue is independent of anorexia-induced muscle mass loss in cancer cachexia. A group of ncxC26 mice was under severe caloric restriction (severeCR), receiving 30% less food intake than a group of cxC26 mice ad lib starting from day 7 post-tumor inoculation. **(a)** Food intake. **(b)** Percentage change in body weight. **(c)** Percentage change of body composition **(d)** Muscle mass. **(e)** Grip strength at the endpoint. **(a-e)** n=6 for ncxC26 ad lib, n=6 for ncxC26 severeCR, and n=5 for cxC26 ad lib. All data are shown as the mean ± s.d. Significance of the differences: ns not significant, *P < 0.05, **P < 0.01, ***P < 0.001, **** P < 0.0001 between groups by one-way ANOVA followed by Tukey post hoc comparison.

### Anorexia can account for important metabolic alterations observed in cancer cachexia

To probe the extent to which anorexia contributes to cancer cachexia-associated metabolic changes, we performed a metabolomics analysis on serum samples collected from the ncxC26 ad lib, isocaloric-fed ncxC26, and isocaloric-fed cxC26 mice. Based on principal component analysis (PCA), isocaloric restriction caused significant serum metabolome changes, demonstrated by the distinct separation between the ncxC26 ad lib cluster and the isocaloric-fed ncxC26 cluster (Fig. 6a). Note that the isocaloric-fed cxC26 cluster diverged from the isocaloric-fed ncxC26 cluster, suggesting the presence of tumor-dependent metabolic effects. We further identified significantly altered metabolites (adjusted p-values < 0.05) across comparisons with isocaloric-fed cxC26 versus ncxC26 ad lib representing the conventional comparison, isocaloric-fed ncxC26 versus ncxC26 ad lib showing the caloric restriction-induced metabolic changes, and isocaloric-fed cxC26 versus isocaloric-fed ncxC26 showing anorexia-independent metabolic changes. As shown in the Venn diagram, 31 out of 51 caloric restriction-induced metabolites overlapped with those identified in the conventional comparison (Fig. 6b), suggesting that metabolomics results on cancer cachexia can be misleading in terms of reflecting metabolic abnormalities if food intake is not controlled for. Only 26 out of 72 significantly altered metabolites from the conventional comparison dataset overlapped with net cancer cachexia-associated metabolic changes (Supplementary Fig. S3). Changes in important metabolic indicators such as glucose and ketone bodies (acetoacetate and 3-hydroxybutyrate) (Fig. 6c) and liver glycogen content (Fig. 6d) are likely consequences of reduced food intake instead of reflecting metabolic abnormalities in cancer cachexia.

**Figure 6.**
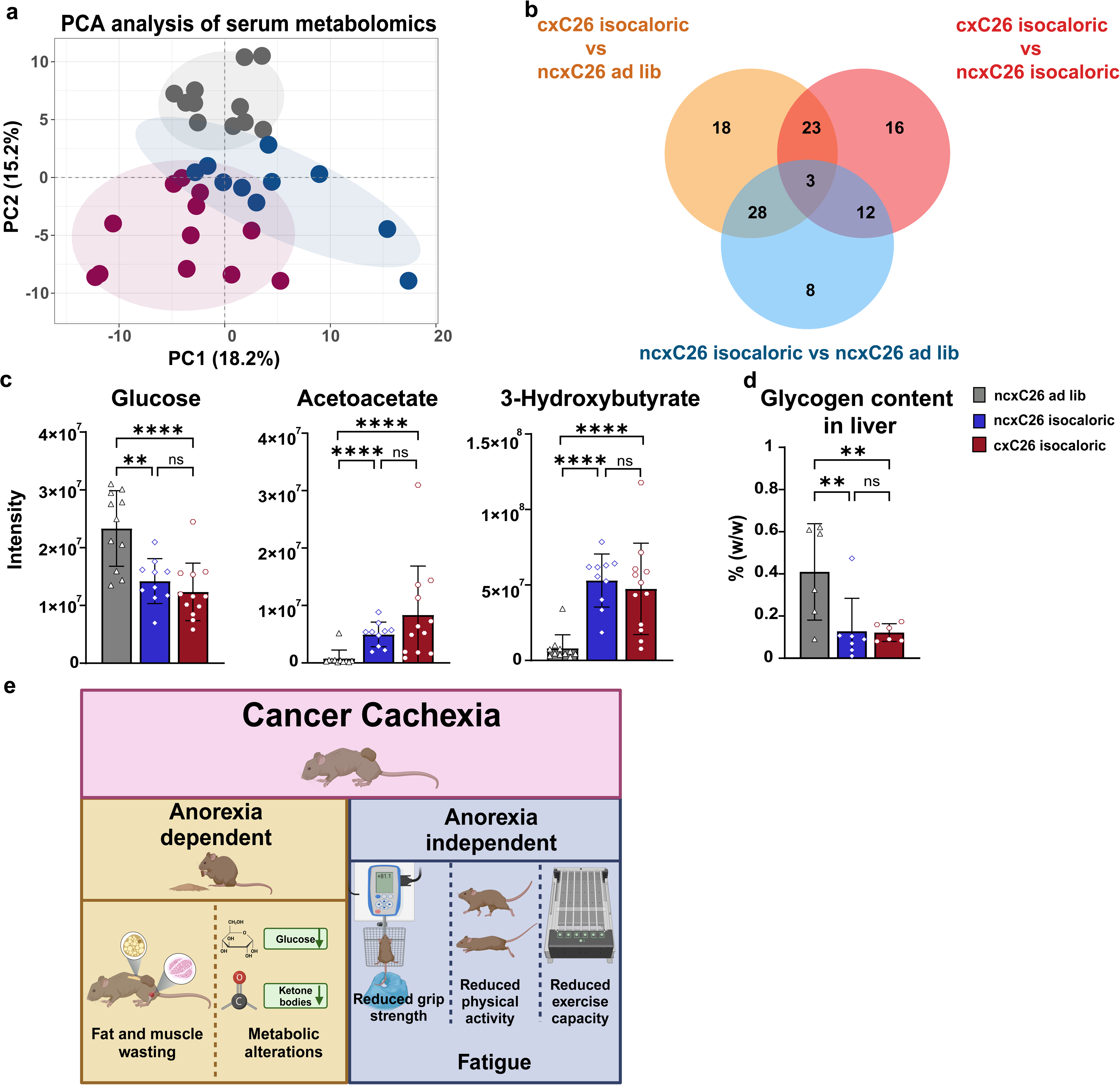
Anorexia accounts for important metabolic alterations observed in cancer cachexia. **(a)** Principle component analysis of the serum metabolomics. **(b)** Venn diagram of the significant metabolites found in different comparisons. **(c)** Serum glucose and ketone bodies, acetoacetate and 3-hydroxybutyrate. **(d)** Liver glycogen content. **(e)** Diagram summarizing the anorexia-dependent and -independent cachectic phenotypes in the C26 model. **(a, c)** n=11 for ncxC26 ad lib, n=10 for isocaloric-fed ncxC26, n=12 for isocaloric-fed cxC26. **(d)** n=6 for ncxC26 ad lib, n=7 for isocaloric-fed ncxC26, n=6 for isocaloric-fed cxC26.

## Discussion

This study demonstrates the importance of separating anorexia-dependent and -independent effects in cancer cachexia research. The findings that key cachectic phenotypes, including body weight loss, mass loss of adipose tissue and muscle, and alterations of key metabolites, are mostly contributable to anorexia in the C26 model inform that a search for new mechanisms for these phenotypes may not be justified in this model. Instead, the finding that fatigue occurs independently of anorexia highlights novel mechanisms that warrant further investigation. The extent to which anorexia contributes to cachectic phenotypes is generally different in different preclinical models and different subtypes of patients. Thus, we propose to make it a standard procedure to get accurate food intake data and perform rigorous control for food intake in studies of cancer cachexia.

Our results caution using the expression of the genes *Murf1* and *Atrogin1* as markers of muscle atrophy in cancer cachexia. Consistently with the muscle mass data, food intake reduction alone induces their expression to the same level as in the cachectic C26 mice. Thus, while these gene markers may reflect muscle mass loss, their use as an indication of active muscle wasting processes is not validated.

It was surprising to find that the loss of adipose tissue and muscle mass was simply due to the food intake reduction in the C26 model. This is because the model has often been used for studying active mechanisms of adipose tissue wasting [14,35] and muscle wasting [36–38]. These studies generally did not check food intake or conduct food intake control, and thus cannot distinguish whether the wasting was primarily due to food intake reduction or some active mechanisms. While we cannot rule out the complete lack of additional mechanisms of tissue wasting, our results indicate a minor role for them in this model.

Our study showed no elevation of energy expenditure in the C26 model. Despite being commonly cited as a contributor to body weight loss in cancer cachexia, energy expenditure elevation has only been observed in limited number of cases, including some patients with lung and pancreatic cancer [39], as well as preclinical models such as Lewis lung cancer (LLC) cachexia model [23] and a skin cancer cachexia model [40].

Our results revealed that fatigue does not necessarily depend on food intake reduction or muscle mass loss in cancer cachexia. With comparable anorexia-induced muscle loss, only the cachectic animals exhibited reductions in grip strength, physical activities, and maximum exercise capacity, while non-cachectic animals remained as competent as the ad lib control. In line with our results, a study using a lung cancer cachexia model compared caloric-restricted mice with cachectic mice and observed similar reductions in fat mass, lean mass, and skeletal muscle mass, but no change of muscle function in the caloric-restricted group [41]. Grip strength has been reported to be preserved [42–44], and exercise endurance has been shown to increase [45,46] in humans after caloric restriction, which aligns with what we found in the calorie-restricted non-cachectic mice. Thus, different from muscle wasting or muscle atrophy, fatigue is a distinct phenotype in cancer cachexia. Fatigue could be due to either muscle function impairment [47,48] or neural dysfunction [49–51], which so far has been understudied in cancer cachexia research and deserves more attention in the field.

By isolating out the effects of reduced food intake, the goal is to simplify the complex problem of cancer cachexia and ultimately establish the causal pathways of cancer cachexia. We do not downplay the role of anorexia in cancer cachexia. Instead, as shown in our results, anorexia plays a dominant role in multiple phenotypes of cachexia and thus deserves intensive study. We simply argue for separate treatments of anorexia-dependent and -independent phenotypes while searching for the underlying mechanisms (Fig. 7). In this structure of causal pathway, a successful strategy for therapeutical development might be to act separately on appetite and physical performance. In line with our view, several clinical trials targeting the ghrelin pathway for increased food take had no effects on hand grip strength and overall survival despite reported improved lean body mass [52–54]. While inhibition of GDF15 signaling was reported to have mitigated not only body weight loss but also physical performance impairment through rescuing appetite in preclinical cancer cachexia models (HT-1080 [55] and TOV21G (ovarian cancer) [56]), the mechanism by which GDF15 impacts physical performance is unclear and the physical performance in a recent clinical trial of an anti-GDF15 antibody had inconsistent dose-dependence [57]. Thus, a future effective treatment is likely a combination of two drugs, targeting anorexia and fatigue.

**Figure 7.**
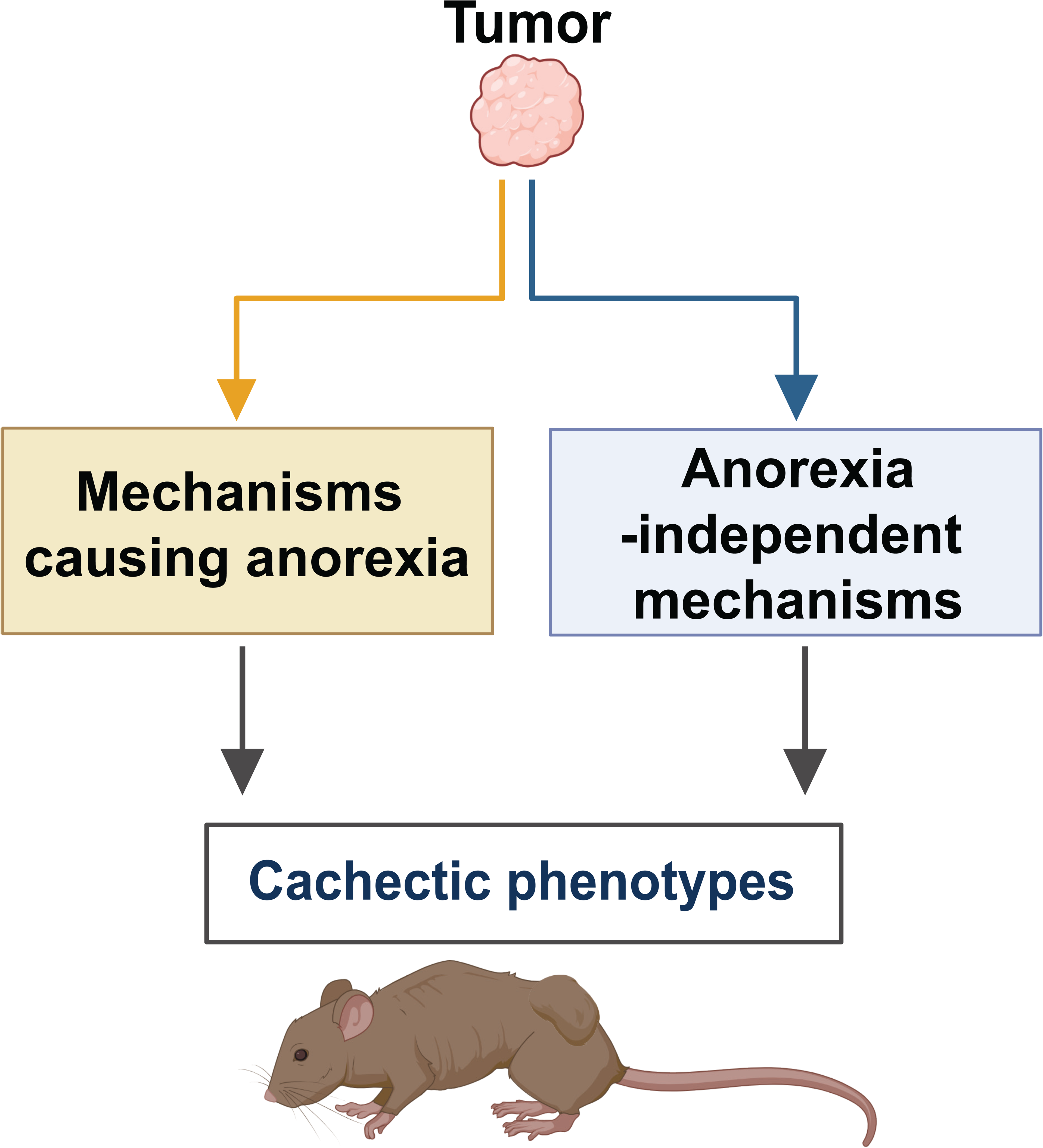
**The causal pathway of cancer cachexia consists of mechanisms causing anorexia and anorexia-independent mechanisms.**

## Methods

### Animals

CD2F1 male mice were purchased from Charles River Laboratories and were housed (4 per cage) under regular light-dark cycles of 12 h with ad lib access to water and food (PicoLab 5053, LabDiet). Prior to tumor implantation, mice were single-housed and acclimated to experimental conditions for 1 week. Tumor implantation was conducted when animals were 12-16 weeks old and reached a stable stage in growth and weight gain. One million of cxC26 or ncxC26 cells per animal were implanted subcutaneously in a 200-µL injection of saline cell suspension into the right flank of mice. The length of the studies was 2 weeks. Animal care and experimental procedures were conducted with the approval of the Institutional Animal Care and Use Committees (IACUC) of Harvard Medical School and Harvard T.H. Chan School of Public Health.

### Cell culture

The cachectic C26 (cxC26) and non-cachectic (ncxC26) cells were kind gifts from Andrea Bonetto [17] and Nicole Beauchemin, respectively. Nicole Beauchemin obtained the ncxC26 cells from Brattain MG, who established the C26 (also named CT26 or colon carcinoma 26) cell line [58]. All the cell lines were maintained using high glucose DMEM (Corning) with 10% fetal bovine serum (FBS, R&D Systems) and 1% penicillin/streptomycin (P/S, Hyclone). Cells were maintained in incubators at 37 °C and 5% CO_2_.

### Food intake and isocaloric feeding

For food intake measurement and during the isocaloric feeding, animals were housed individually on alpha pad bedding (6” × 10”, scottPharm solution) with virgin kraft crinkled paper strips as nesting material (Enviro-Dri™, scottPharm solution) (See Extended Data Fig. 2a). Food pellets were weighed and placed each day on the lid. The mass of food pellets on the lid from the previous day was checked and recorded the next day. Any visible food debris (Extended Data Fig. 2a) on the bedding was collected, weighed, and discarded. The food surplus would be the sum of pellets on the lid and the bedding. Daily food intake was calculated as the weight difference in food pellets between 2 consecutive days.

For the isocaloric feeding, the mice in all three groups were body weight-matched. A group of ncxC26 tumor-bearing mice were on ad lib as a control. From day 6 to day 12 after tumor injection, the cxC26 mice and another group of ncxC26 mice were fed the same amount of food daily following the food intake of the cxC26 mice we measured in Fig. 1a. Food debris on the bedding will be checked and weighed daily and returned to the mice. On day 12, any mice with more than 1g of food (about 5% of the total food intake of the isocaloric feeding) left would be excluded from the experiment.

### Body weight and body composition assessment

Body weight was measured every other day following implantation (approximately 11 am and 1 pm). Whole-body composition (fat mass and lean mass) analysis was performed using an EchoMRI-100 (EchoMRI LLC) on each animal at day 0 and day 12 prior to their euthanasia. Each mouse was individually placed in a specialized plastic tube for mild conscious restraint and inserted into a magnetic resonance imager (MRI) to measure body composition. Each mouse took approximately 5 min to complete the procedure. Mice were then returned to their home cage.

### Metabolic cage analyses

Mice were individually housed at 22 °C and subjected to indirect calorimetry between day 5 and day 12 after tumor inoculation under a 12 h light-dark cycle using the Promethion Metabolic Cage system (Sable Systems, USA). During this period, spontaneous activity, and volume of oxygen consumed and carbon dioxide produced were measured. This data allows us to calculate energy expenditure (EE) and the respiratory exchange ratio (RER). Fold changes in daily locomotion and cage distance were calculated by comparing values from day 12 to day 0.

### Fecal energy determination

Feces produced 24 hours before the endpoint were collected and stored at -80 °C. The feces were dehydrated at 60 °C for 48 hours in an Eppendorf tube with lid open. Then, the calorie content of the dry feces was determined by using Bomb Calorimetry (Parr Oxygen Bomb). Wet and dry weights were measured for later calculations of total fecal production (g/day) and fecal caloric content (cal/g).

### Muscle mass and grip strength

At the end of the study, animals were euthanized, and gastrocnemius, soleus, tibialis anterior (TA), and extensor digitorum longus (EDL) were isolated and weighed. To measure grip strength, mice were allowed to grab the metal grid and then were pulled backward by an experimenter until the grasp was released. The peak tension was measured using a Grip Strength meter (Bioseb).

### Maximum exercise capacity test

Mice were acclimated to a motorized treadmill for three consecutive days from day 4-6 before the maximal exercise capacity test. Shocking grids with frequency set at 2 Hz and intensity at 1.22 mA were located at the end of the treadmill to force the mice to run at their maximum capacity. The treadmill was set at 10° incline. The running test was conducted at the end of day 12. Animals ran at 6 m/min for the first 2 min, at 10 m/min for the following 8 min, followed by 2 m/min increases every 5 min until exhaustion, defined by the inability to remain on the treadmill for >5 s.

### RNA extraction and RT-qPCR

Fresh gastrocnemius samples were isolated from ncxC26 ad lib, isocaloric-fed ncxC26, and isocaloric-fed cxC26 tumor-bearing mice and immediately snap-frozen using liquid nitrogen. Total RNA was extracted using the Trizol reagent (Invitrogen, #15596026) from the 20-30 mg of gastrocnemius and TA muscle tissues, following the manufacturer’s protocol. RNA concentration was determined using NanoDrop One (Thermofisher Scientific). One microgram of purified RNA was converted to cDNA using the iScript Reverse Transcription Supermix for RT-qPCR (Bio-Rad, #1708840). The diluted cDNA was used for RT-qPCR using SsoAdvanced Universal SYBR Green Supermix (Bio-Rad, #1725270) with the CFX384 Real-Time PCR detection System (Bio-Rad). The following primer sequences were used to determine the Atrogin-1 and MuRF1 mRNA expression. Cyclophilin was used as a housekeeping gene for normalization.

**Table 1.**
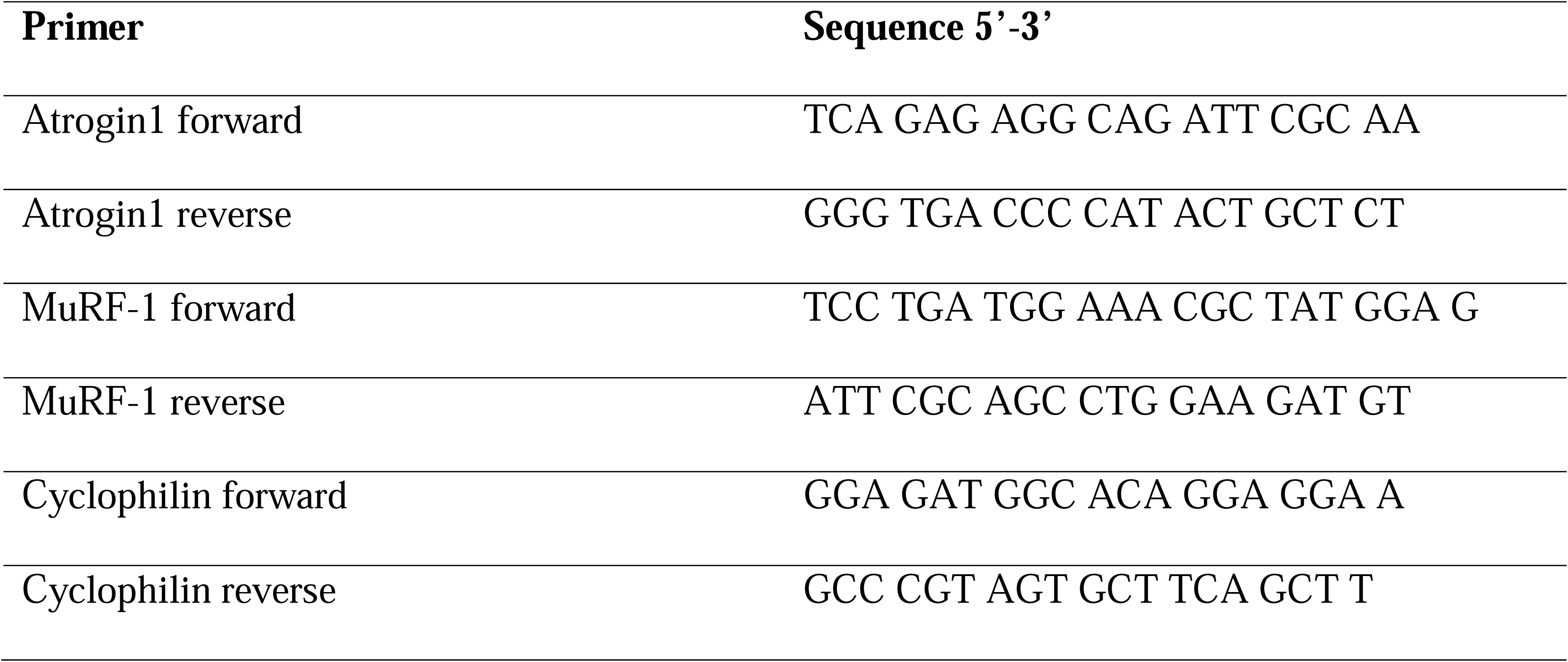
Primer sequences.

### Measurement of glycogen in tissues

Mice were fasted for 7-8 hours before the tissue collection. They were euthanized by cervical dislocations, and tissues (liver, gastrocnemius, and soleus) were immediately dissected, clamped with a pre-chilled Wollenberger clamp, and rapidly submerged in liquid nitrogen for snap-1freezing. Frozen tissues were then ground under cryogenic condition using a CryoMill (Retsch). For glycogen quantification, 10 – 20 µg of ground tissue were used in the Glycogen-Glo assay kit (Promega, #J5051), following the manufacturer’s instructions.

### Serum metabolomics

Serum metabolites were extracted using extraction buffer (acetonitrile: methanol: water, 4: 4: 2, *v: v: v*) at a volume-to-volume ratio of 1:20 (μL to μL), followed immediately by vortexing for 10 seconds. The mixture was then incubated at 4 °C for 10 min before centrifugation at 16,000 g at 4 °C for 10 min. The supernatant was transferred to an LC-MS vial and analyzed on the LC-MS. A pooled quality control (QC) sample was prepared by mixing 20 μL from each experimental sample.

Metabolite extracts were analyzed using a quadrupole-orbitrap mass spectrometer coupled with hydrophilic interaction chromatography (HILIC). Chromatographic separation was achieved on an XBridge BEH Amide XP Column (2.5 µm, 2.1 mm × 150 mm) with a guard column (2.5 µm, 2.1 mm × 5 mm) (Waters, Milford, MA). For the gradient, mobile phase A was water: acetonitrile 95:5, and mobile phase B was water: acetonitrile 20:80, both phases containing 10 mM ammonium acetate and 10 mM ammonium hydroxide. The linear elution gradient was: 0 ∼ 3 min, 100% B; 3.2 ∼ 6.2 min, 90% B; 6.5. ∼ 10.5 min, 80% B; 10.7 ∼ 13.5 min, 70% B; 13.7 ∼ 16 min, 45% B; and 16.5 ∼ 22 min, 100% B, with a flow rate of 0.3 mL/ min. The autosampler was at 4°C. The column temperature was at 30°C. The injection volume was 5 µL. Needle wash was applied between samples using acetonitrile: methanol: water at 4: 4: 2 (*v: v: v*). For mass spectrometry, a Q Exactive HF (Thermo Fisher Scientific, San Jose, CA) mass spectrometer was used. For MS1 acquisition, the MS scanned from 70 to 1000 m/z with switching polarity at a resolution of 120,000 for all experimental samples. The relevant parameters were: sheath gas, 40; auxiliary gas, 10; sweep gas, 2; spray voltage, 3.5 kV; capillary temperature, 300 °C; S-lens, 45. The resolution was set at 120,000 (at m/z 200). Maximum injection time (max IT) was set at 500 ms, and automatic gain control (AGC) was set at 3 × 10^6^.

MS1 raw data files were converted into mzxML using msconvert and imported to EI-Maven (Elucidata, Cambridge, MA) for targeted metabolomics. Metabolites were identified using an in-house library based on accurate mass and retention time.

### Statistical analysis

All statistical analyses for murine data were performed in GraphPad Prism 10 or R studio. Quantitative data are reported as mean with standard deviation (s.d.). Three-group comparisons were analyzed using one-way ANOVA followed by Tukey post hoc. Metabolites with adjusted p-values less than 0.05 are considered significant. For all analyses, a p-value of <0.05 was considered significant (*p < 0.05, **p < 0.01, and ***p < 0.001, ****p < 0.0001).

---

**Supplementary Figure S1. Pair-feeding recapitulates the body weight loss and body composition of the cachectic C26 mice ad lib.** CD2F1 male mice were inoculated with 1×10^6^ cxC26 or ncxC26 cells. A group of ncxC26 mice were pair-fed to match the food intake of cxC26 mice ad lib. **(a)** Percentage change of body weight. **(b)** Body composition. **(c)** Tumor mass. **(a-c)** n=8 for ncxC26 ad lib, n=8 for pair-fed ncxC26, and n=8 for cxC26 ad lib.

**Supplementary Figure S2. Cage setup for food intake measurement and masticated food debris in cages of cxC26 and ncxC26 mice. (a)** Cage setup. Mice were single-housed with alpha paper bedding and paper strips as nesting material. **(b)** Mass of food debris in cages of ncxC26 and cxC26 mice (n=8 per group). The cxC26 mice produced significantly more food debris than the ncxC26 mice. The mass of food debris should be considered when calculating the daily food intake.

**Supplementary Figure S3. Volcano plot for serum metabolites between isocaloric-fed cxC26 and ncxC26 groups.** Metabolites with adjusted p-values < 0.05 were considered as significant changes, with elevated metabolites shown in red and decreased ones in green. N=10 for isocaloric-fed ncxC26, n=12 for isocaloric-fed cxC26.

## Acknowledgements

We would like to thank Dr. Andrea Bonetto (cxC26) and Dr. Nicole Beauchemin (ncxC26) for graciously providing the cell lines used for our studies. We are grateful to Dr. Tobias Janowitz, Dr. Eileen White, and Dr. Marcus Goncalves for their critical reading and valuable suggestions of the manuscript. We thank Karen E. Inouye for the technical support with the metabolic cages and EchoMRI, and Guangru Jiang for the help in fecal energy excretion measurement. This work was delivered as part of the CANCAN team supported by the Cancer Grand Challenges partnership funded by Cancer Research UK (CGCATF-2021/100034) and the National Cancer Institute (OT2CA278654).

## Author contributions

Yanshan Liang: Conceptualization; Resources; Investigation; Data curation; Formal analysis; Visualization; Methodology; Writing—original draft; Project administration; Writing—review and editing. Young-Yon Kwon: Conceptualization; Resources; Investigation; Writing—original draft; Writing—review and editing. Sheng Hui: Conceptualization; Supervision; Funding acquisition; Investigation; Writing original draft; Project administration; Writing—review and editing.

