## Supplementary figures and images for "Separating anorexia -dependent and -independent effects in cancer cachexia"

### Figure S1-3

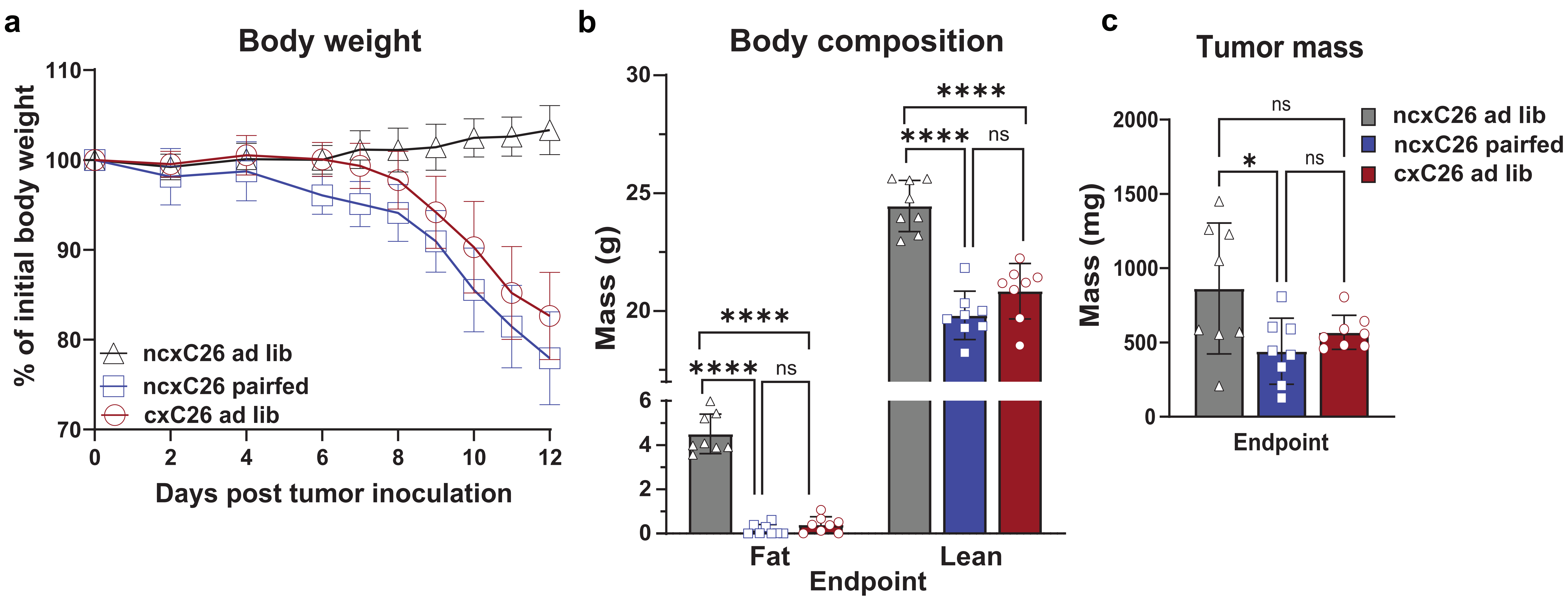

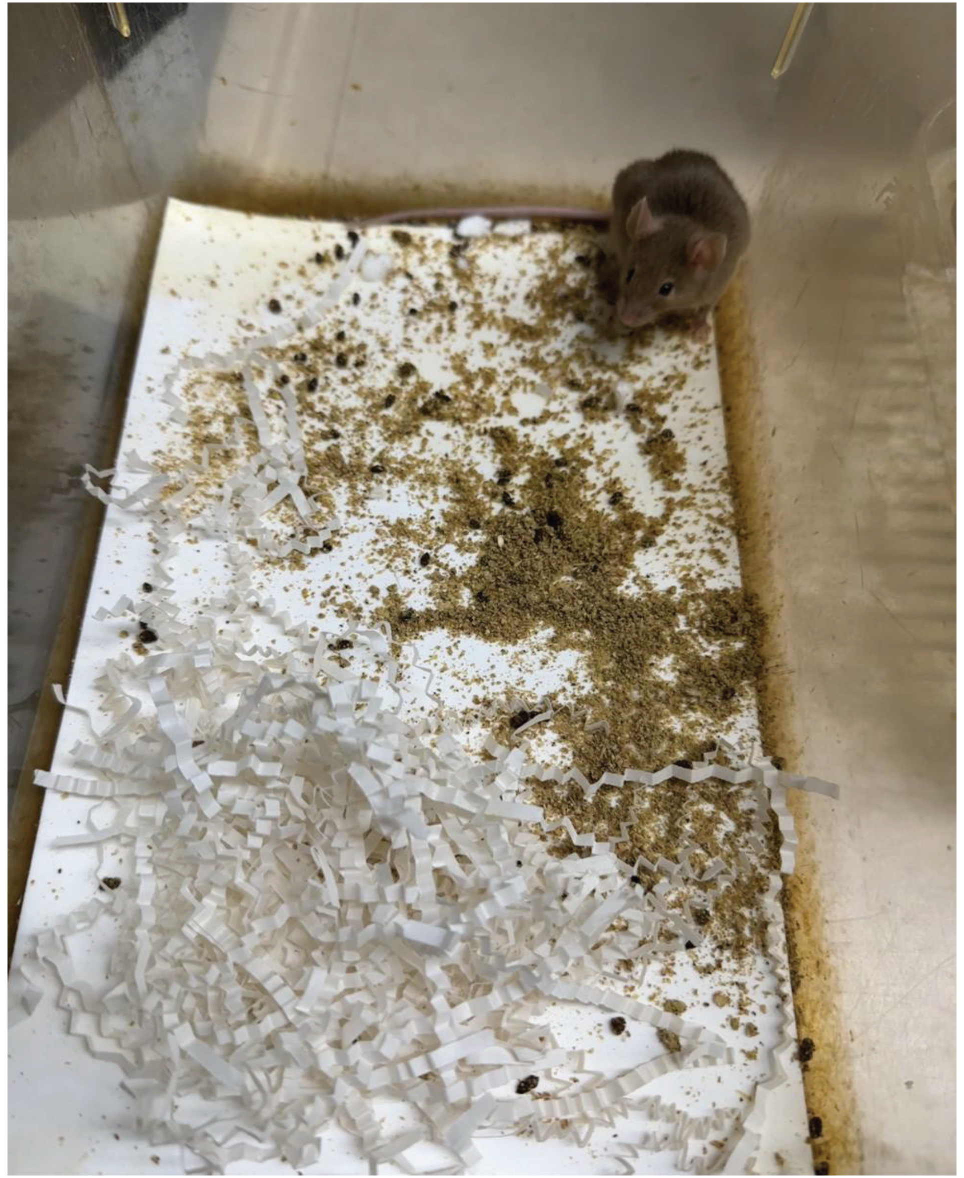

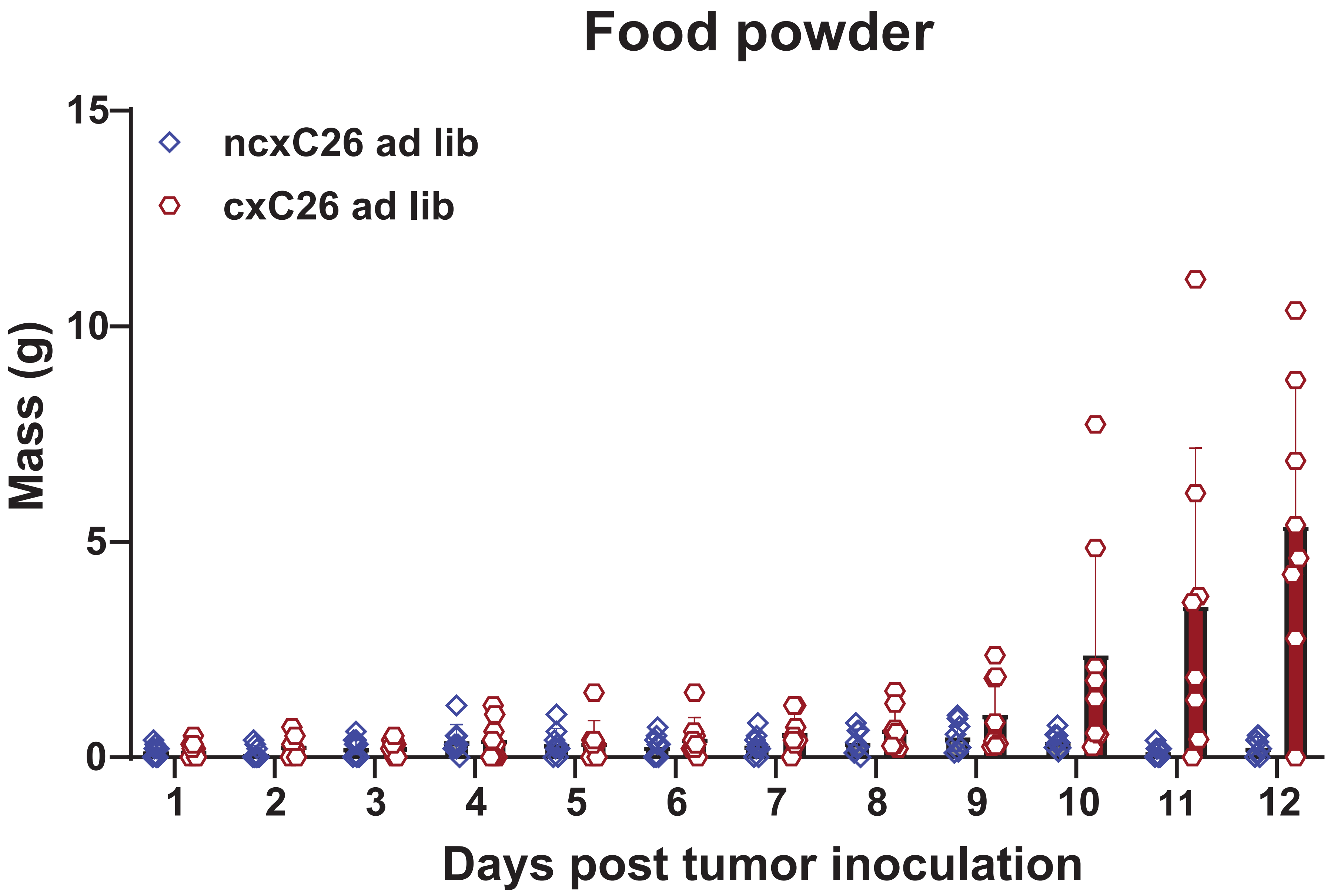

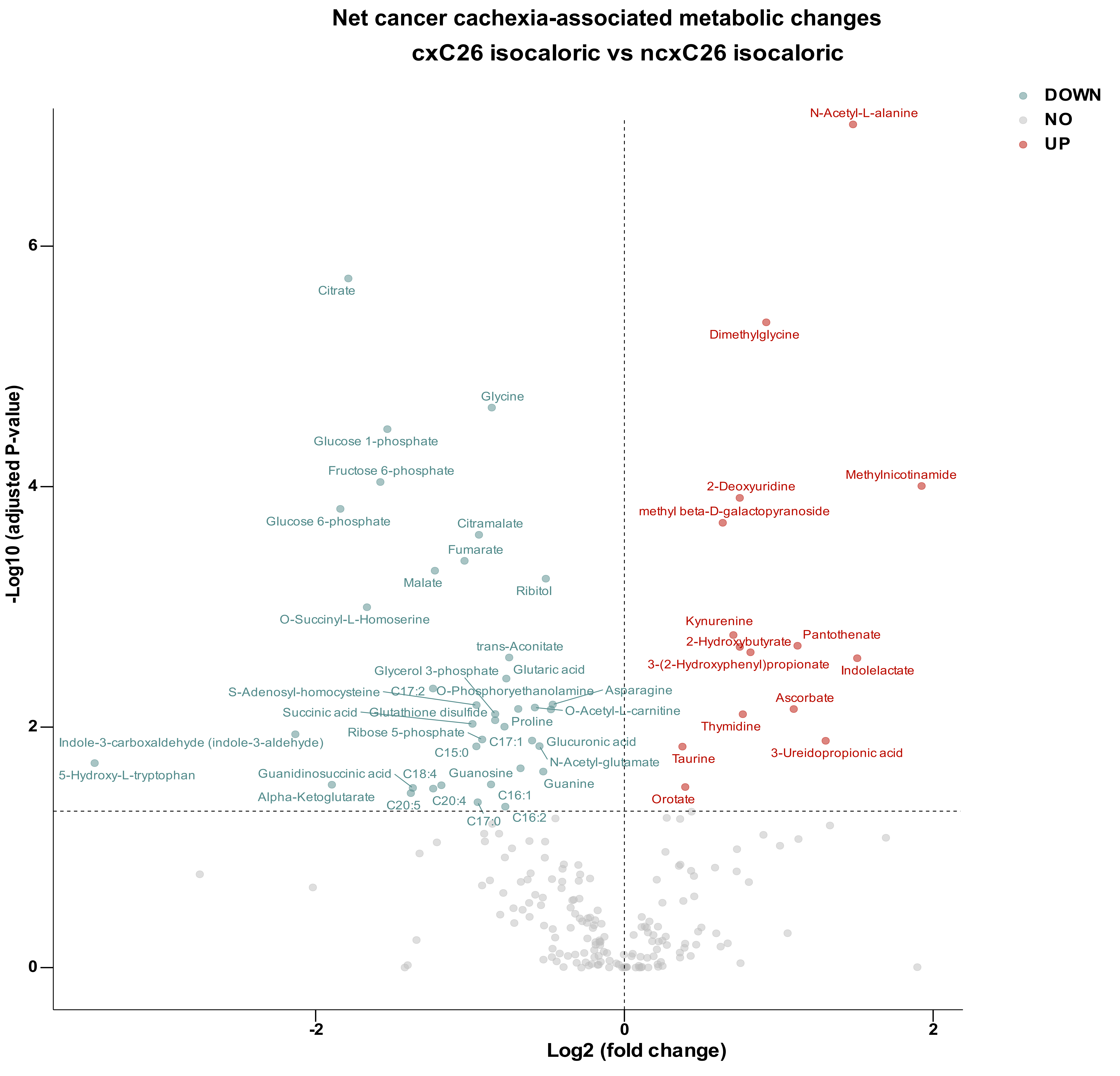
